# The metalloproteinase Papp-aa functions as a molecular switch linking IGF signaling to adaptive epithelial growth

**DOI:** 10.1101/792978

**Authors:** Chengdong Liu, Shuang Li, Pernille Rimmer Noer, Kasper Kjaer-Sorensen, Caihuan Ke, Claus Oxvig, Cunming Duan

## Abstract

Human patients carrying inactivating mutations in the pregnancy-associated plasma protein-a2 (PAPP-A2) gene display short status and lower bone mineral density. The underlying mechanisms are not well understood. Using a zebrafish model, here we report a [Ca^2+^]-dependent mechanism by which Papp-aa regulates bone calcification via promoting Ca^2+^-transporting epithelial cell (ionocyte) reactivation. Ionocyte, normally quiescent, re-enter the cell cycle in response to low [Ca^2+^] stress. Deletion of Papp-aa abolished ionocyte reactivation and resulted in a complete lack of calcified bone. Re-expression of Papp-aa, but not its active site mutant, rescued ionocyte reactivation. Inhibition of Papp-aa activity pharmacologically or by overexpressing STC1 or STC2 impaired ionocyte reactivation. Loss of Papp-aa expression or activity resulted in diminished IGF1 receptor-mediated Akt-Tor signaling activity in ionocytes and expression of a constitutively active Akt rescued ionocyte reactivation. Biochemically, Papp-aa cleaved Igfbp5a, a high-affinity IGF binding protein specifically expressed in ionocytes. Under normal [Ca2+] conditions, the Papp-aa-mediated Igfbp5a proteolysis was suppressed and IGFs sequestered in the IGF/Igfbp5a complex. Forced release of IGFs from the complex was sufficient to activate the IGF-Akt-Tor signaling and promote ionocyte reactivation. These findings suggest that Papp-aa functions as a [Ca^2+^]-regulated molecular switch linking IGF signaling to adaptive epithelial growth and bone calcification.

## Introduction

Pregnancy-associated plasma protein-a (PAPP-A) and PAPP-A2 belong to the conserved pappalysin protein family (Boldt et al., 2001; Oxvig, 2015; Conover and Oxvig, 2018). PAPP-A was first discovered in the plasma of pregnant women in the 1970s. Decades of studies suggest that PAPP-A and PAPP-A2 are zinc metalloproteinases specifically cleaving insulin-like growth factor binding proteins (IGFBPs) (Laursen et al., 2007; Oxvig, 2015; Conover and Oxvig, 2018). IGFBPs are a family of secreted proteins that bind IGF ligands with high affinity and regulates their availability to the IGF1 receptor (Baxter, 2014; Clemmons, 2018; Allard and Duan, 2018). In vitro studies suggested that PAPP-A is tethered to the cell-surface and it cleaves IGFBP4, IGFBP5, and to a lesser degree IGFBP2, while PAPP-A2 is secreted and mainly cleaves IGFBP3 and IGFBP5 (Oxvig, 2015). Papp-a knockout mice showed a 40% reduction in body size (Conover et al., 2004), a phenotype similar to those of the IGF1 and IGF2 mutant mice (Baker et al., 1993; Liu et al., 1993). Papp-a2 knockout mice showed a modest decrease in body size (Christians et al., 2006; 2013). These findings have led to the proposal that PAPP-A and PAPP-A2 promote somatic growth by increasing bioavailable IGFs (Oxvig, 2015; Fujimoto et al., 2017; Conover and Oxvig, 2018). This notion is further supported by recent clinical studies showing that human patients carrying inactivating mutations in the PAPP-A2 gene showed progressive growth failure (Dauber et al., 2016; Fujimoto et al., 2017). In addition to reduced body height, these patients also had lower bone mineral density (Dauber et al., 2016; Fujimoto et al., 2017). Likewise, the Papp-a mutant mice had delayed appearance of ossification centers and reduced calcified bone mass (Conover et al., 2004). The Papp-a2 mutant mice had sex-and age-specific defects in bone structure and mineral density (Christians et al., 2019). These observations suggest a role of pappalysin metalloproteinases in bone mineralization, but causal evidence is still lacking and the underlying mechanisms are not well understood.

We have recently found that genetic deletion of the major Papp-a/Papp-a2 substrate IGF binding protein 5a (Igfbp5a) resulted in greatly reduced calcified bone mass even though *igfbp5a* is not expressed in skeletal tissues in zebrafish (Liu et al., 2018). Zebrafish *igfbp5a* is specifically expressed in a population Ca^2+^-transporting epithelial cells located in the yolk sac epidermis (Dai et al., 2014; Liu et al., 2017). These cells, known as ionocytes or NaR cells, are functionally similar to human intestinal epithelial cells (Hwang, 2009; Lin and Hwang, 2016). A hallmark of NaR cells and human intestinal epithelial cells is the expression of Trpv6/TRPV6, a conserved calcium channel that constitutes the first and rate-limiting step in the transcellular Ca^2+^ transport pathway (Hoenderop et al., 2005; Pan et al., 2005; Dai et al., 2014; Xin et al., 2019). NaR cells take up Ca^2+^ from the surrounding aquatic habitats to maintain body Ca^2+^ homeostasis. These cells, normally non-dividing and quiescent, rapidly exit the quiescent state and proliferate in response to low [Ca^2+^] stress (Dai et al., 2014; Liu et al., 2017). Genetic deletion of *igfbp5a* blunted the low [Ca^2+^] stress-induced IGF signaling and NaR cell proliferation (Liu et al., 2018). This is likely an adaptive response, allowing fish to take up adequate Ca^2+^ to maintain body Ca^2+^ homeostasis and survive under low [Ca^2+^] conditions. Although *igfbp5a*^−/−^ fish did not show overt phenotypes when raised in high [Ca^2+^] solutions, they died prematurely in normal and lower [Ca^2+^] media (Liu et al., 2018), suggesting a [Ca^2+^]-dependent mechanism(s) exists.

In this study, we show that Papp-aa is highly expressed in NaR cells and genetic deletion of Papp-aa resulted in a complete lack of calcified bone and abolished low [Ca^2+^] stress-induced NaR cell reactivation and proliferation. We provide several lines of evidence showing that Papp-aa acts by cleaving Igfbp5a and activating IGF signaling in NaR cells to promote the quiescence-proliferation transition in response to low [Ca^2+^] stress. Interestingly, the Papp-aa proteolytic activity is suppressed under normal [Ca^2+^] conditions, suggesting that Papp-aa-mediated Igfbp5a proteolysis acts as a [Ca^2+^]-regulated switch linking local IGF signaling to adaptive epithelial proliferation and bone calcification.

## Results

### Papp-aa is highly expressed in NaR cells and is indispensable in NaR cell reactivation

Zebrafish has three genes belonging to the pappalysin family, *papp-aa*, *papp-ab*, and *papp-a2* (Kjaer-Sorensen et al., 2013, 2014; Wolman et al., 2015). To date, published whole mount *in situ* hybridization data showed that *papp-aa* mRNA is expressed in various neural tissues, *papp-ab* mRNA in developing myotomes and brain (Kjaer-Sorensen et al., 2013;Wolman et al., 2015; Miller et al., 2018; Alassaf et al., 2019), and *papp-a2* in the notochord and brain (Kjaer-Sorensen et al., 2014). Because NaR cells are located on the yolk sac skin, they are more sensitive to permeabilization by protease K treatment, a key step in the whole mount *in situ* hybridization procedure. Thus, to assess pappalysin expression, NaR cells were isolated by FACS from *Tg(igfbp5a:GFP)* fish, in which NaR cells are labeled by GFP expression (Liu et al., 2017) and analyzed. The mRNA levels of *papp-aa* were 2-fold higher than those of *papp-ab* and *papp-a2* (Fig. 1A). Low [Ca^2+^] stress treatment had no effect on their levels (Fig. 1A). We also compared the mRNA levels of *papp-aa* in NaR cells to those in non-GFP labeled cells. The levels of *papp-aa* mRNA in NaR cells were approximately 10-fold greater (Fig. 1B). In comparison, the *papp-ab* mRNA levels were similar between NaR cells and other cells (Fig. 1C). Next, whole mount *in situ* hybridization was performed after optimizing the permeabilization condition. In agreement with previous reports (Wolman et al., 2015; Miller et al., 2018), strong *papp-aa* mRNA signals were detected in the brain (Fig. 1D). Meanwhile, *papp-aa* mRNA signal was detected in cells in the yolk sac region at 3dpf or later (Fig. 1D). Double label staining showed that *papp-aa* mRNA is indeed expressed in NaR cells (Fig. 1E).

**Figure 1.**
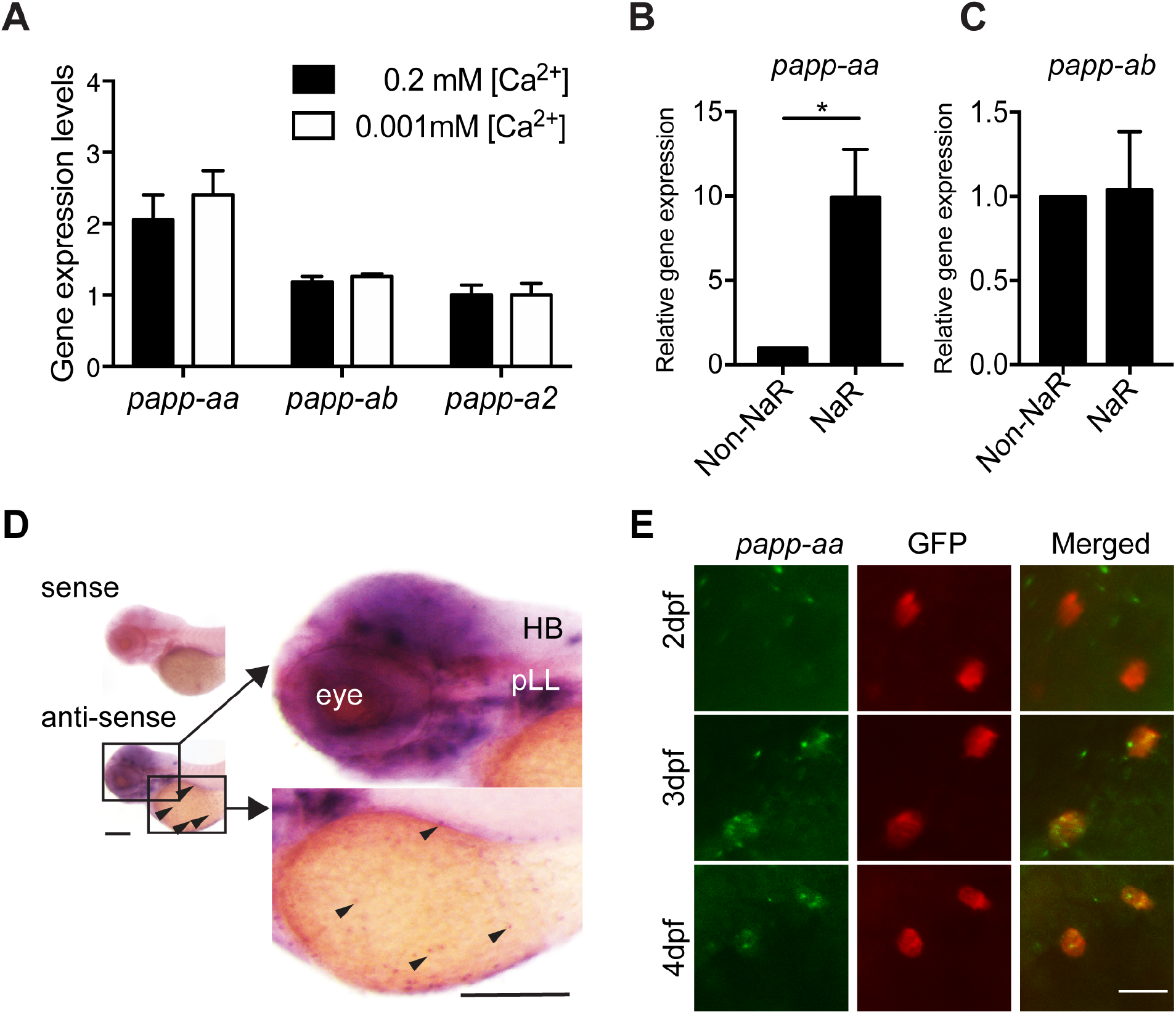
Papp-aa is highly expressed in NaR cells. (**A**) *Tg(igfbp5a:GFP)* fish were raised in E3 embryo medium to 3 days post fertilization (dpf) and transferred to embryo media containing the indicated [Ca^2+^]. Eighteen hours later, NaR cells were isolated by FACS. The levels of *papp-aa*, *papp-ab*, and *papp-a2* mRNA were measured and shown. Data shown are Mean ± SEM, n= 4. (**B-C**) NaR cells and other cells in 4 dpf *Tg(igfbp5a:GFP)* larvae were separated by FACS. The levels of *papp-aa* (**B**) and *papp-ab* (**C**) mRNA were measured and shown. *, *P* < 0.05 by unpaired two-tailed t test. n = 3. (**D**) Whole mount *in situ* hybridization analysis of *papp-aa* mRNA in 3 dpf larvae. HB, hindbrain. pLL, posterior lateral line ganglion. Arrowheads indicate *papp-aa* mRNA signal in the yolk sac region. A sense cRNA probe was used as a negative control. Scale bar = 0.2 mm. (**E**) *Tg(igfbp5a:GFP)* fish of the indicated stages were analyzed by double label staining. Scale bar = 20 µm.

The possible role of Papp-aa in NaR cell reactivation was investigated by double blind experiments (Fig. 2A) using the progeny of *papp-aa*^*+/−*^ fish intercrosses (Wolman et al., 2015, referred as *pappaa^p170^*^/+^). Under normal [Ca^2+^] condition, NaR cell number and density among these 3 different genotype groups were similar (Fig. 2B and 2C). Low [Ca^2+^] stress treatment resulted in a 4-fold, highly significant increase in NaR cell number (labeled by *igfbp5a* mRNA) in wild-type and heterozygous fish, whereas this increase was abolished in *papp-aa^−/−^* mutant fish (Fig. 2B and 2C). Similar results were obtained using *trpv6* mRNA as a NaR cell marker (Supplemental Fig. S1A). NaR cells in wild-type larvae had markedly enlarged apical opening under low [Ca^2+^] stress treatment (Fig. 2D). Although the functional significance of this morphological change is not clear, it was not abolished in *papp-aa^−/−^* mutant fish (Fig. 2D). Next, we examined NaR cell reactivation using the progeny of *papp-aa*^*+/−*^;*Tg(igfbp5a:GFP)* fish intercrosses. Deletion of Papp-aa completely abolished NaR cell reactivation (Fig. 2E and 2F).

**Figure 2.**
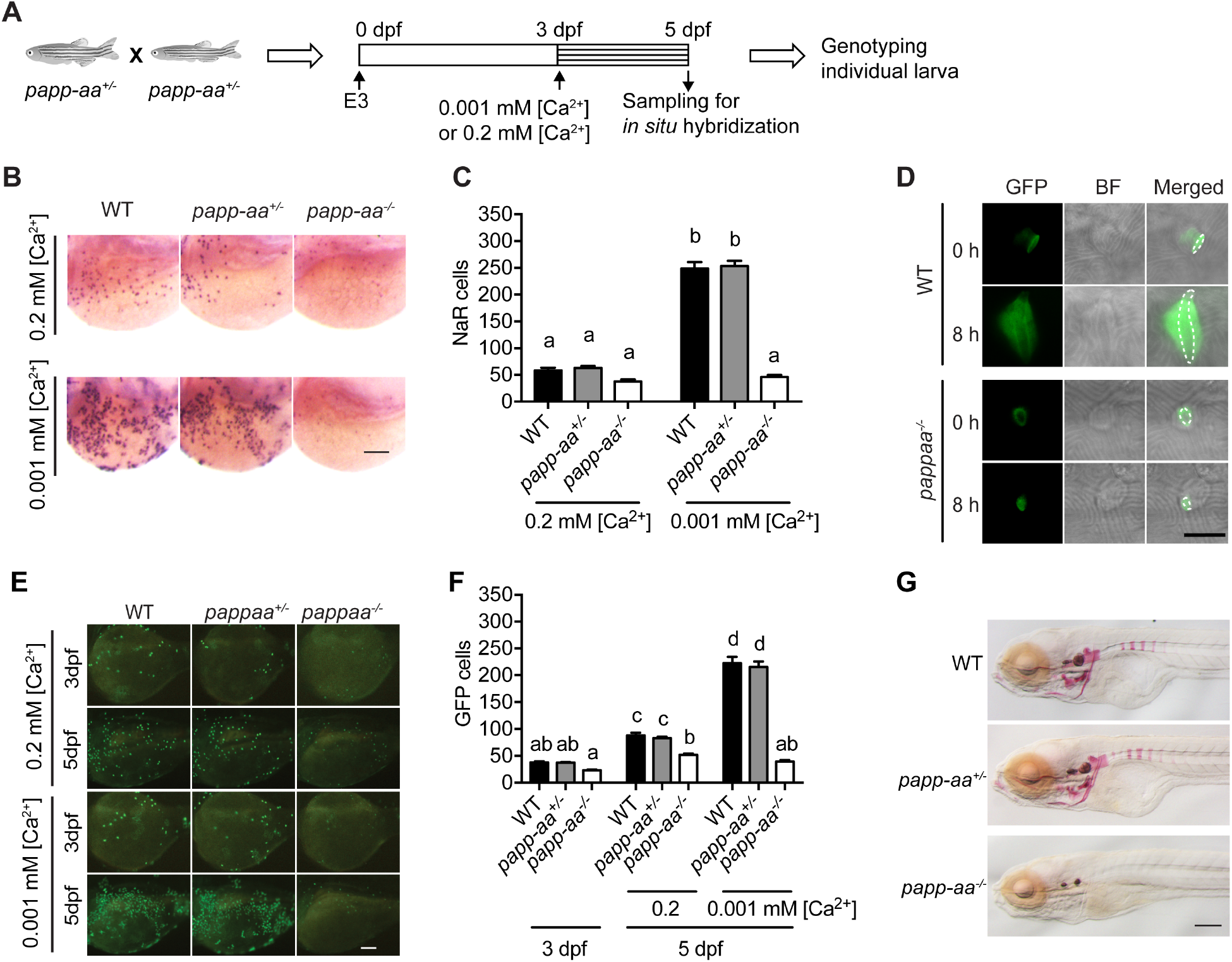
Genetic deletion of *papp-aa* impairs NaR cell reactivation and bone calcification. **(A)** Diagram of the experimental design. Progeny of *papp-aa*^+/−^ fish intercrosses were raised in E3 embryo medium to 3 dpf and transferred to the low [Ca^2+^] (0.001 mM) or normal [Ca^2+^] (0.2 mM) embryo medium. Two days later, NaR cells in each fish were labeled by *igfbp5a* mRNA expression and quantified. These fish were genotyped individually afterwards. (**B-C**) Progeny of *papp-aa*^*+/−*^ intercrosses were treated as described in (**A**). Representative images are shown in **(B)** and quantified data in (**C**). Scale bar = 0.1 mm. n = 10~30 fish/group. In this and all subsequent figures, data shown are Mean ± SEM. Different letters indicate significant differences among groups by one-way ANOVA followed by Tukey’s multiple comparison test (*P* < 0.05). (**D**) Progeny of *papp-aa^+/−^;Tg(igfbp5a:GFP)* fish intercrosses were raised in E3 medium to 3 dpf. NaR cells were imaged before and 8 hours after the low [Ca^2+^] treatment. The apical opening was marked by dotted lines. Scale bar = 10 µm. (**E-F**) Progeny of *papp-aa*^*+/−*^;*Tg(igfbp5a:GFP)* fish were subjected to the low [Ca^2+^] stress test described in (**A**). The number of GFP-expressing NaR cells in each larva was quantified. The larvae were genotyped individually subsequently. Representative images are shown in (**E**) and quantified data in (**F**). Scale bar = 0.1 mm. n = 16~82 fish/group. (**G**) Fish of the indicated genotypes were raised in E3 embryo medium to 7 dpf and stained with Alizarin Red. Scale bar = 0.2 mm.

To test whether the action of Papp-aa is restricted to NaR cells, we examined H^+^-ATPase-rich (HR) cells and Na^+^/Cl^−^ cotransporter (NCC) cells, two other types of epithelial cells located in the larval yolk sac responsible for H^+^ secretion/Na^+^ uptake/NH^+^ excretion and Na^+^uptake/Cl^−^ uptake, respectively (Hwang, 2009). Deletion of Papp-aa did not change HR and NCC cell numbers (Supplemental Fig. S1B and C). No difference was observed between mutant fish and siblings in term of gross appearance, growth rate, and developmental timing (Supplemental Fig. S2). As reported by Wolman et al. (2015), all mutant fish died around 2 weeks. Alizarin red staining indicated a complete lack of calcified bone in *papp-aa^−/−^* fish (Fig. 2G), suggesting they suffered from Ca^2+^ deficiency. Together, these data suggest that Papp-aa plays an indispensable role in regulating body Ca^2+^ balance, bone calcification, and NaR cell reactivation in response to low [Ca^2+^] stress.

### Papp-aa is an Igfbp5a proteinase and endogenous Papp-aa proteinase activity is critical for quiescence reactivation

To determine whether Papp-aa expression is sufficient in inducing NaR cell reactivation, Papp-aa was randomly expressed in NaR cells in *pappaa^−/^*^−^; *Tg(igfbp5a:GFP*) fish using a Tol2 transposon BAC-mediated genetic mosaic assay (Liu et al., 2018). Re-introduction of Papp-aa restored low [Ca^2+^] stress-induced NaR cell reactivation (Fig. 3A and 3B). To test whether this action of Papp-aa requires its proteinase activity, a conserved glutamate residue (E501) of the active site known to be critical for human PAPP-A proteinase activity (Boldt et al., 2001) was changed to alanine. As expected, Papp-aa E501A showed no proteolytic activity (Fig. 3C). When the E501A mutant was expressed randomly in NaR cells in *pappaa*^*−/−*^; *Tg(igfbp5a:GFP*) using the genetic mosaic assay, it had no effect on NaR cell reactivation (Fig. 3A and 3B).

**Figure 3.**
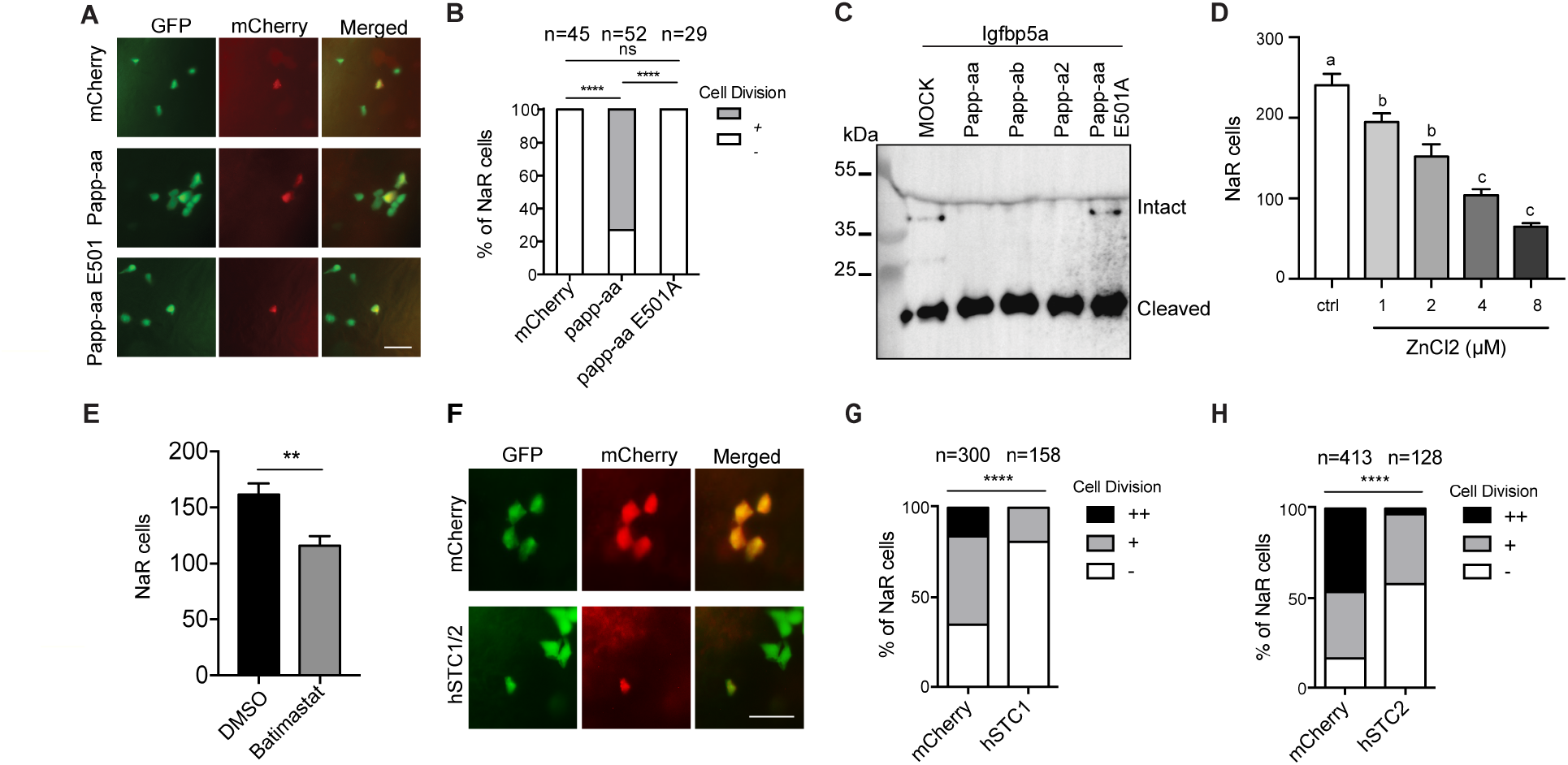
Papp-aa proteinase activity is critical in NaR cell reactivation. **(A-B**) Progeny of *papp-aa*^*+/−*^;*Tg(igfbp5a:GFP)* intercrosses were injected with *BAC(igfbp5a:mCherry)* containing the indicated gene. They were subjected to the low [Ca^2+^] stress test described in Fig. 2A. Papp-aa-IRES-mCherry, Papp-aa E501A-IRES-mCherry, or mCherry expressing NaR cells were detected by GFP and mCherry expression. NaR cells expressing mCherry or Papp-aa-mCherry (yellow, double labeled by GFP and mCherry) were scored following a published scoring system (Liu et al., 2018). Representative images are shown in (**A**) and quantified data in (**B**). Scale bar = 50 µm. +, one cell division, 0, no division. ****, P< 0.0001 by Chi-square test. The total cell number is shown above the bar. (**C**) Conditioned media collected from HEK293 cells co-transfected with Igfbp5a and the indicated plasmid were analyzed by Western blotting. Intact and cleaved Igfbp5a bands were indicated. (**D-E**) *Tg(igfbp5a:GFP)* fish were transferred to the low [Ca^2+^] medium containing 0-8 µM ZnCl_2_ (**D**) or 200 µM Batimastat at 3 dpf (**E**). After two days of treatment, NaR cells were quantified and shown. n = 18~25 fish/group. **, P< 0.001 by unpaired two-tailed t test. (**F-H**) *Tg(igfbp5a:GFP)* embryos were injected with *BAC(igfbp5a:mCherry, BAC(igfbp5a:hSTC1-IRES-mCherry)* (**G**) or *BAC(igfbp5a:hSTC2-IRES-mCherry*) (**H**). They were raised and subjected to the low [Ca^2+^] stress test described in Fig. 2A. NaR cells expressing mCherry or human STC (yellow, double labeled by GFP and mCherry) were scored following a published scoring system (Liu et al., 2018). Representative images are shown in (**F**) and quantified results in (**G and H**). ++, two cell division, +, one cell division, 0, no division during the experiment. ****, *P*< 0.0001, Chi-square test. Total cell number is shown above the bar.

Next, *Tg(igfbp5a:GFP)* embryos were treated with ZnCl_2_ or batimastat, two metzincin metalloproteinase inhibitors (Tallant et al., 2006). Both compounds inhibited low [Ca^2+^] stress-induced NaR cell reactivation (Fig. 3D and 3E). Human stanniocalcin (STC) 1 and 2 can directly bind and inhibit PAPP-A proteinase activity (Kløverpris et al., 2015; Jepsen et al., 2015). When they were each targeted expressed in NaR cells randomly in *Tg(igfbp5a:GFP)* embryos using the genetic mosaic assay (Liu et al., 2018), the low [Ca^2+^] stress-induced reactivation was significantly inhibited in TSC1 or STC2 expressing NaR cells (Fig. 3F-H). These results suggest that the presence of proteolytically active Papp-aa is critical for NaR cell quiescence reactivation.

Zebrafish has two paralogous *igfbp5* genes, but no *igfbp4* (Dai et al., 2010; Allard and Duan, 2018). To test whether Papp-aa directly cleaves Igfbp5a and to investigate the relationship among multiple Igfbp5 and pappalysins, in vitro proteinase assays were carried out. Both Papp-aa and Papp-ab cleaved Igfbp5a efficiently (Fig. 4A), whereas Papp-a2 showed little activity (Fig. 4A). In comparison, Papp-aa, Papp-ab, and Papp-a2 all cleaved the paralogous Igfbp5b (Fig. 4A), but only Papp-a2 was able to cleave the closely related Igfbp3 (Fig. 4A). Human IGFBP5 mutated at position K128 (K128A) resists cleavage by PAPP-A (Laursen et al., 2001; 2002). We therefore mutated two basic residues located in the corresponding positions of Igfbp5a, K147 and K148, to alanine. The Igfbp5a K148A mutant showed a modest reduction in Papp-aa-mediated cleavage and a marked reduction in Papp-ab-mediated cleavage (Fig. 4B and 4C). Changing K147 to alanine had little effect (Fig. 4B and 4C). Mutational analysis of the paralogous Igfbp5b showed that the K145A and K144A/K145K mutants were resistant to Papp-aa and Papp-ab with the double mutant having a stronger effect (Fig. 4D and 4E). In comparison, Igfbp5b K145A mutant had little effect (Fig. 4D and 4E). Thus, Papp-aa and Papp-ab, but not Papp-a2, can cleave Igfbp5a and Igfbp5b and cleavage specificity depends on residues which are conserved between human and zebrafish proteins.

**Figure 4.**
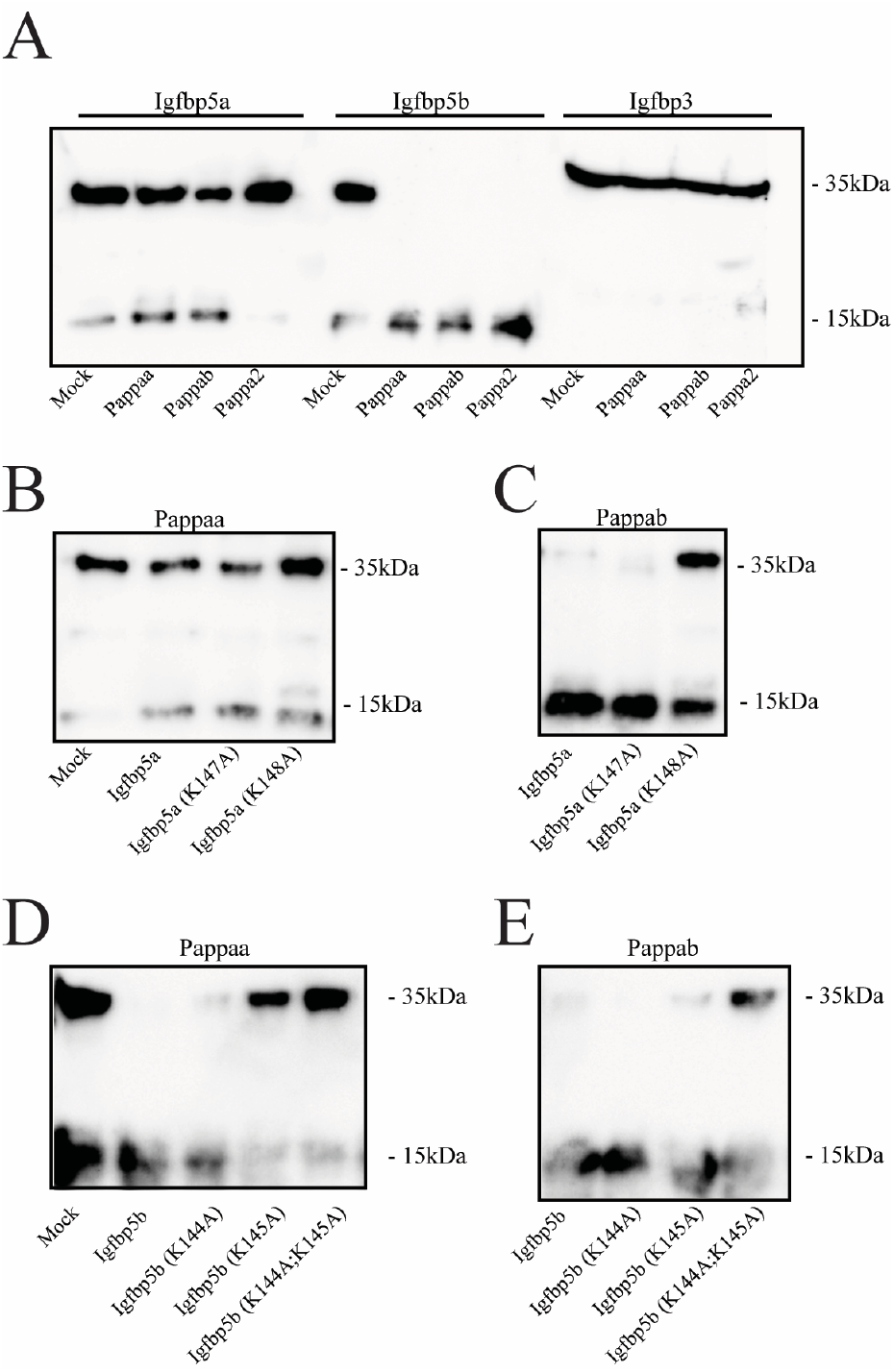
Papp-aa is an Igfbp5 proteinase. (**A**) Igfbp5a, Igfbp5b, or Igfbp3 was incubated with the indicated Papp-a protein. For the Papp-a2 and Igfbp3 group, 100 nM IGF-II was added. After 24 h, the protease reaction was stopped and analyzed by Western blotting. Intact and cleaved Igfbp bands are indicated. **(B-C)** Igfbp5a and the indicated Igfbp5a mutant protein were incubated with Papp-aa (**B**) or Papp-ab (**C**) for 24 h. The proteinase reaction was stopped and analyzed by Western blotting. **(D-E)** Igfbp5b and the indicated Igfbp5b mutant protein were incubated with conditioned media collected from HEK293 cells transfected with Papp-aa (**D**) and Papp-ab (**E**) for 24 h. The protease reaction was stopped and analyzed by Western blotting.

### Papp-aa promotes NaR cell reactivation via proteolytic cleavage of Igfbp5a and increasing IGF signaling activity

In agreement with previous reports (Dai et al., 2014; Liu et al., 2018), low [Ca^2+^] stress treatment activated NaR cell Akt and Tor activity in wild-type fish (Fig. 5A and 5B; Supplemental Fig. S3). This induction was impaired in *papp-aa*^*−/−*^ fish but not in the heterozygous fish (Fig. 5A and 5B; Supplemental Fig. S3). Treatment of wild-type zebrafish larvae with ZnCl_2_ and batimastat inhibited the low [Ca^2+^] stress-induced Akt and Tor activity (Fig. 5C and 5D), indicating that Papp-aa may promote NaR cell reactivation by activating Akt-Tor signaling. If this were correct, then constitutive activation of this signaling pathway should rescue NaR cell reactivation in *papp-aa*^−/−^ fish. Indeed, genetic mosaic assays showed that targeted expressing of myr-Akt, a constitutively active Akt, in NaR cells in *papp-aa*^*−/−*^ fish, restored their reactivation in response to low [Ca^2+^] stress (Fig. 5E and 5F).

**Fig. 5.**
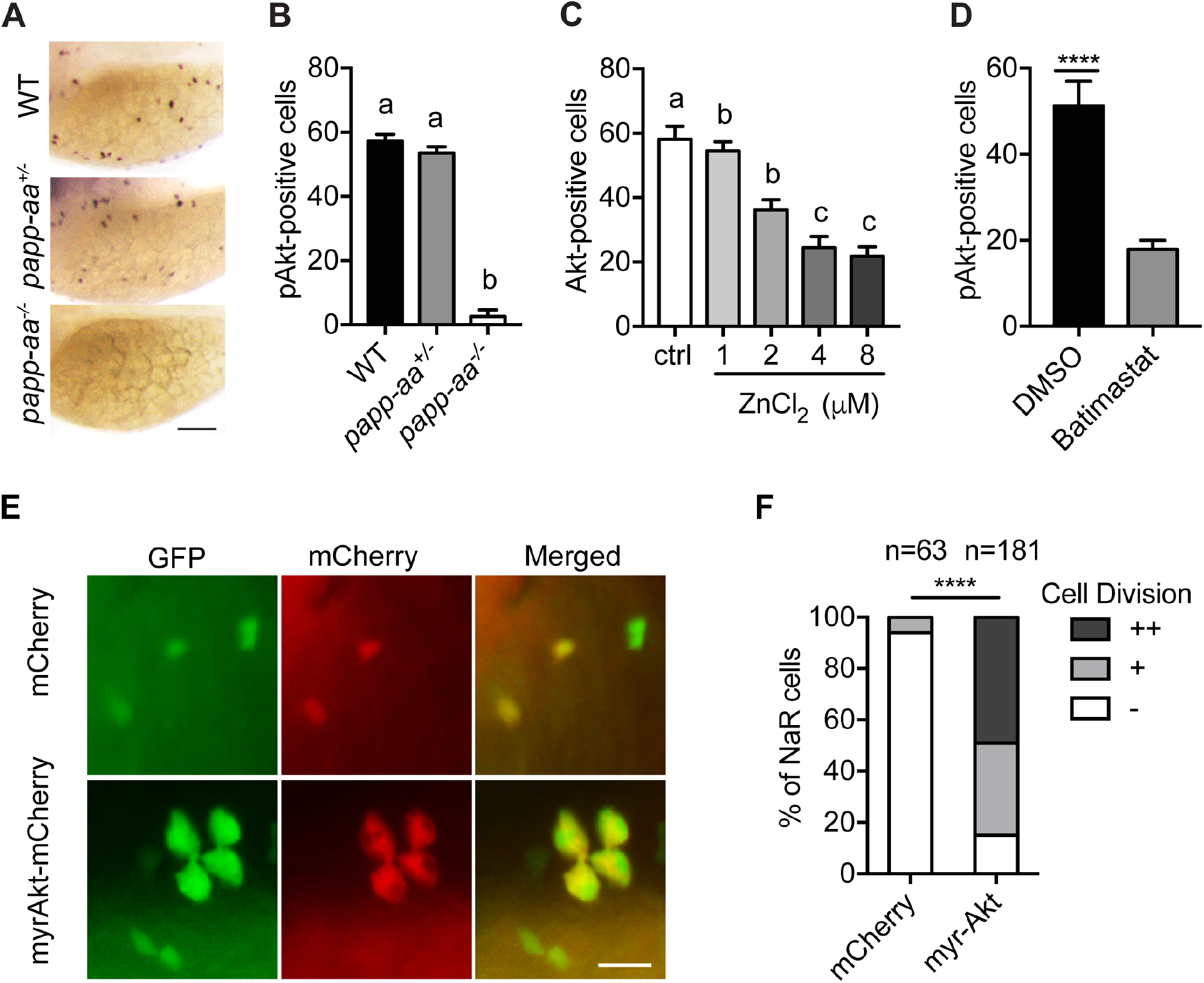
Papp-aa acts by regulating IGF-Akt-Tor signaling in NaR cells. (**A-B**) Zebrafish embryos of the indicated genotypes were transferred to the low [Ca^2+^] medium at 3 dpf. One day later, they were fixed and stained for phospho-Akt. Representative images are shown in (**A**) and quantitative results in (**B**). Scale bar = 0.1mm. n = 25-58 fish/group. **(C-D**) *Tg(igfbp5a:GFP)* fish were transferred to the low [Ca^2+^] medium 0-8 µM ZnCl_2_ (**C**) or 200 µM batimastat (**D**) at 3 dpf. After one day treatment, they were analyzed by immunostaining for phospho-Akt. n = 18~24 fish/group. ****, p<0.0001, unpaired two-tailed t test. (**E and F**) Progeny of *pappaa^+/−^;Tg(igfbp5a:GFP)* intercrosses were injected with *BAC(igfbp5a:mCherry)* or *BAC(igfbp5a:myr-Akt-mCherry)*. They were raised and subjected to the low [Ca^2+^] stress test described in Fig. 2A. NaR cells expressing mCherry or myr-Akt were scored as described in Fig. 3. Representative images are shown in (**E**) and quantified data in (**F**). ++, two cell division, +, one cell division, 0, no division during the experiment. Scale bar = 20 µm. ****, *P*< 0.0001 by Chi-square test. Total number of cells is shown above the bar.

Because Papp-aa increases IGF signaling in NaR cells only under low [Ca^2+^] stress, we speculated that Papp-aa activity is suppressed in normal [Ca^2+^] and IGF ligands are sequestered in the Igfbp5a/IGF complex. If this were correct, then forced release of IGFs from the Igfbp5a/IGF complex should activates the IGF signaling and increase NaR cell reactivation under normal [Ca^2+^] conditions. This idea was tested using NBI-31772, an aptamer that can displace and release IGF from the IGF/IGFBP complex (Chen et al., 2001). Addition of NBI-31772 promoted NaR cell reactivation under normal [Ca^2+^] in a concentration-dependent manner (Fig. 6A and 6B), showing that latent IGF is present and that the limiting step in NaR cell reactivation under normal [Ca^2+^] is the release of bioavailable IGFs. In agreement with this idea, NBI-31772 treatment significantly increased the levels of phospho-Akt and phospho-pS6 activity in NaR cells in an IGF1 receptor dependent manner (Fig. 6C and 6D). Moreover, the effect of NBI-31772 in reactivating NaR cells was abolished by adding the IGF1 receptor inhibitor, BMS-754807, PI3 kinase inhibitor wortmannin, Akt inhibitor MK2260, and Tor inhibitor rapamycin (Fig. 6E; Supplemental Fig. S4B). NBI-31772 treatment did not change the Phospho-Erk levels (Supplemental Fig. S4) and the MEK inhibitor treatment did not affect NBI-31772 treatment-induced NaR cell proliferation (Supplemental Fig. S4B).

**Figure 6.**
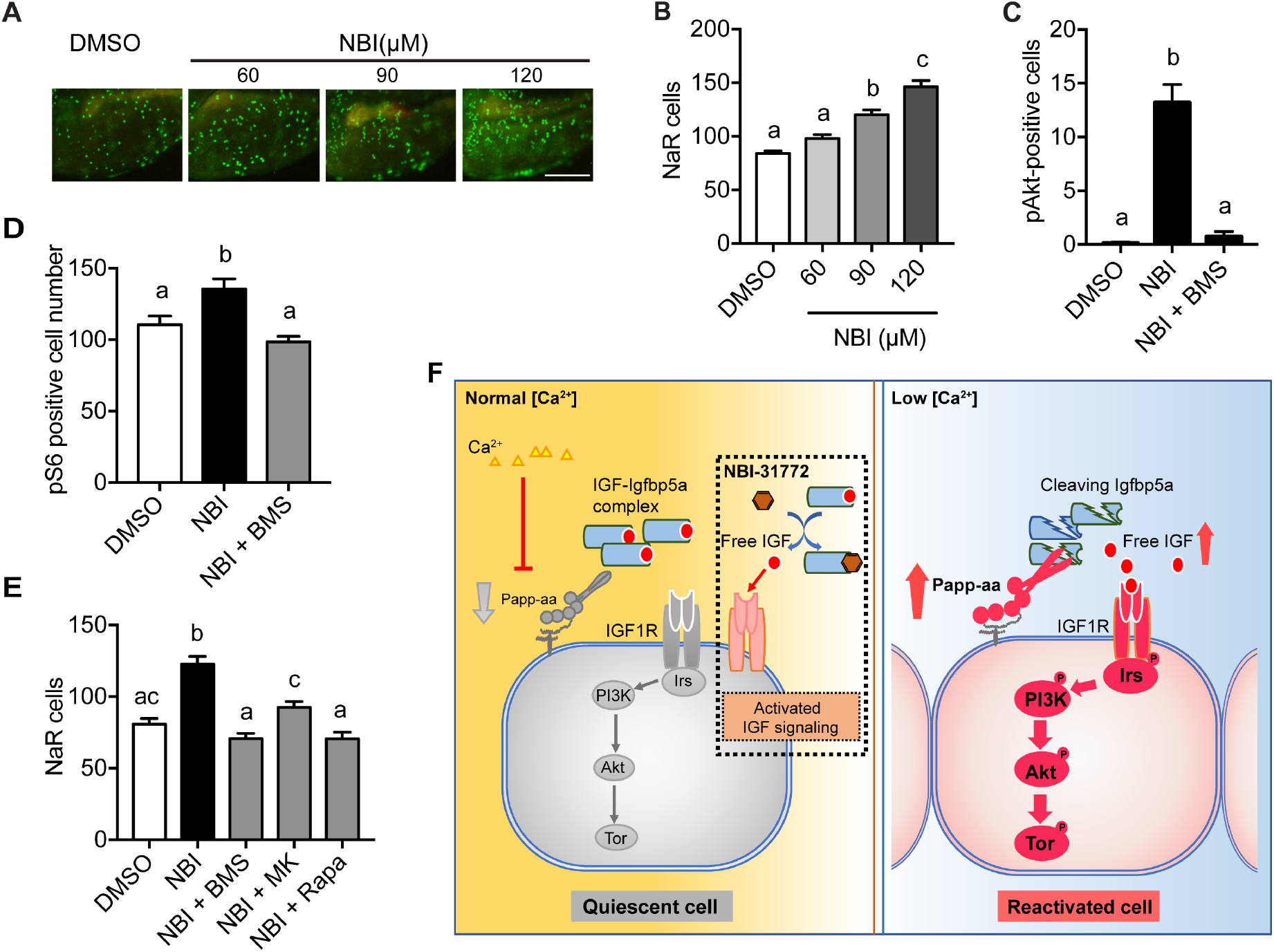
Disruption the IGF/Igfbp complex activates IGF-Akt-Tor signaling and promotes NaR cell reactivation. (**A-B**) *Tg(igfbp5a:GFP)* fish were transferred to normal [Ca^2+^] medium containing the indicated doses of NBI-31772 at 3 dpf. Two days later, NaR cells were quantified. Representative images are shown in (**A**) and quantified data in (**B**). Scale bar = 0.2 mm. n = 21~27 fish/group. (**C**) Wild-type fish were treated with 90 µM NBI-31772 with or without 0.3 µM BMS-754807 from 3 to 4 dpf. The number of cells positive for phosphorylated Akt staining were quantified and shown. n = 9~23 fish/group. (**D**) Larvae treated as described in **(C)** were stained for phosphor S6 and quantified. n = 14~15 fish/group. (**E**) *Tg(igfbp5a:GFP)* fish were treated with NBI-31772 (90 µM) together with BMS-754807 (0.3 µM), MK2206 (8 µM), or Rapamycin (5 µM) from 3 to 5 dpf. NaR cell number was quantified and shown. n =10~24 fish/group. (**F**) Proposed model of Papp-aa function as a [Ca^2+^]-regulated molecular switch linking IGF signaling to reactivation of quiescent epithelial cells. Under normal [Ca^2+^] conditions, Papp-a proteolysis activity is inhibited. Igfbp5a is intact and it binds to IGFs and prevents their binding to the IGF1 receptor. Under low [Ca^2+^] conditions, Papp-a activity is increased. This leads to increased proteolytic cleavage of Igfbp5a and increased liberation of IGFs from the Igfbp5a/IGF complex. This in turn activates IGF-1 receptor-mediated PI3 kinase-Akt-Tor signaling and promotes ionocyte reactivation.

## Discussion

In this study, we uncover a new mechanism linking Papp-aa-mediated Igfbp5a proteolysis to bone calcification and body Ca^2+^ balance and show that Papp-aa acts by reactivating quiescent Ca^2+^-transporting epithelial cells in a [Ca^2+^]-dependent manner. We have recently shown that Igfbp5a is critical in activating IGF signaling and promoting adaptive epithelial growth by reactivating NaR cells in response to low [Ca^2+^] stress (Liu et al., 2018). IGFBP5/Igfbp5a has been shown to inhibit and/or potentiate IGF actions in mammalian and zebrafish cells (Zheng et al., 1998; Ren et al., 2008; Dai et al., 2010; Liu et al., 2018). How the same protein exerts these opposing biological effects is not completely understood, but the importance of IGFBP5 proteinases has been suggested (Zheng et al., 1998; Oxvig, 2015; Clemmons, 2018; Allard and Duan, 2018). Using an optimized *in situ* hybridization protocol and qPCR analysis of FACS-isolated NaR cells, we found that Papp-aa is highly expressed in NaR cells. NaR cells, while largely quiescent under normal [Ca^2+^] conditions, re-enter the cell cycle upon low [Ca^2+^] stress. This adaptive response requires the expression and the proteolytic activity of Papp-aa. Deletion of Papp-aa abolished low [Ca^2+^] stress-induced NaR cell reactivation. Alizarin red staining results showed that *papp-aa*^*−/−*^ fish lacked calcified bone completely. The lack of bone calcification and impaired NaR cell reactivation suggested that *papp-aa*^*−/−*^ fish suffered from Ca^2+^ deficiency, which may explain the premature death of *papp-aa^−/−^* fish. The action of Papp-aa in NaR cells clearly requires its proteinase activity. Reintroduction of wild-type Papp-aa, but not an enzyme inactivated mutant, reactivated NaR cells in the *papp-aa*^*−/−*^ mutant fish. Treatment of wild-type fish with two distinct metalloproteinase inhibitors impaired NaR cell reactivation. Likewise, targeted expression of STC1 and STC2 in NaR cells, two specific PAPP-A inhibitory proteins, blocked low [Ca^2+^] stress-induced NaR cell reactivation.

We provide several lines of evidence showing that Papp-aa regulates the NaR cell quiescence exit decision by activating local IGF signaling. First, the impaired NaR cell reactivation in *papp-aa*^*−/−*^ fish was accompanied with reduced IGF1 receptor-mediated Akt-Tor activity in NaR cells. Second, inhibition of Papp-aa proteolytic activity resulted in reduced IGF-Akt-Tor activity in NaR cells. Third, expression of a constitutively active Akt rescued NaR cell reactivation. Moreover, Papp-aa cleaved Igfbp5a, while Papp-a2 had no such activity. Although Papp-ab can also cleave Igfbp5a in vitro, its expression is relatively low in NaR cells. Finally, the Papp-aa action in NaR cells was blocked by inhibiting the IGF1 receptor, Akt, and Tor. Interestingly, the action of Papp-aa is manifested only under low [Ca^2+^] stress. Papp-aa proteolytic activity is suppressed under normal and physiological conditions because disrupting the IGF/IGFBP complex in fish under normal Ca^2+^ conditions increased Akt and Tor activity and promoted NaR cell reactivation. Our findings therefore suggest that that Papp-aa functions as a [Ca^2+^]-regulated molecular switch. Under normal [Ca^2+^] conditions, Papp-aa proteolytic activity is suppressed. Igfbp5a is intact and IGFs are sequestered in the Igfbp5a/IGF complex. The IGF1 receptor-mediated Akt-Tor activity in NaR cells is off. Under low [Ca^2+^] conditions, Papp-a activity increases and this leads to increased Igfbp5a proteolysis and increased release of bioavailable IGFs from the complex. This activates the IGF1 receptor-mediated signaling and promotes NaR cell reactivation (Fig. 6F).

Since low [Ca^2+^] stress treatment did not change *papp-aa* mRNA levels (this study) or *igfbp5* mRNA levels (Liu et al., 2017) in wild-type fish, and because disruption of the Igfbp/IGF complex was sufficient to activate IGF signaling and induce NaR cell reactivation under normal [Ca^2+^] conditions, we speculate that a [Ca^2+^]-regulated post-transcriptional mechanism(s) exists that suppresses Papp-aa proteolytic activity under normal [Ca^2+^] conditions. Human PAPP-A contains three Lin-12/Notch repeat (LNR) modules (Boldt et al., 2004). LNR modules were initially discovered in the Notch receptor family (Lovendahl et al., 2018). All Notch receptors have 3 LNRs arranged in tandem and they each bind one Ca^2+^. These LNR modules are involved in the ligand-induced cleavage and activation of the Notch receptor (Vardar et al., 2003; Gordon et al., 2007). Interestingly, calcium depletion or deletion of the LNR modules leads to premature activation of Notch signaling (Rand et al., 2000). In the case of PAPP-A, LNR1 and 2 are located in the proteolytic domain, while LNR3 is located in the C-terminus, separated by 1000 residues from LNR 1 and 2 (Boldt et al., 2004). All PAPP-A LNR modules have been shown to bind Ca^2+^ (Boldt et al., 2004). Although separated in the primary structure of PAPP-A, the LNR modules together form a Ca^2+^-dependent substrate binding exosite (Weyer et al. 2007; Mikkelsen et al., 2008). All of these LNR modules are conserved in zebrafish Papp-aa. Whether one or more of these LNR modules are involved in the external [Ca^2+^]-regulated Papp-aa activation is unclear at present.

Alternatively, Papp-aa activity might be suppressed by inhibitory proteins. In vitro studies showed that mammalian PAPP-A proteolytic activity is inhibited by STC1 and STC2 via direct protein-protein binding (Kløverpris et al., 2015; Jepsen et al., 2015). In this study, we showed that overexpression of either STC1 or STC2 in NaR cells impaired Papp-aa-induced NaR cell reactivation. Stc protein was originally discovered in bony fish as a hypocalcemia hormone (Pang, 1973). Recent studies showed that multiple STC genes are present in human and other vertebrate genomes (Yeung et al., 2012). In zebrafish, the mRNA levels of *stc1* but not the other *stc* genes are significantly higher in fish raised in high [Ca^2+^] medium compared to those kept in low [Ca^2+^] medium (Chou et al., 2015). Forced expression and morpholino-based knockdown of Stc1 altered NaR cell density and Ca^2+^ uptake (Tseng et al., 2009), although these manipulations have also changed HR and NCC cell density and the uptake of other ions (Chou et al., 2015). Future studies will clarify whether endogenous Stc1 and/or other Stc proteins are involved in the [Ca^2+^]-dependent activation of Papp-aa.

In summary, the results of this study have uncovered a novel function of Papp-aa in reactivating quiescent epithelial cells and maintaining body Ca^2+^ balance and bone Ca^2+^ content. Papp-aa promotes NaR cells reactivation by cleaving local Igfbp5a and activating the IGF1 receptor-mediated PI3 kinase-Akt-Tor signaling in these cells. These findings not only provide new insights into the physiological functions of the pappalysin family zinc metalloproteinases, but also have important implications in the cellular quiescence regulation. Reactivation of quiescent cells is critical for tissue repair, regeneration, and adult stem cell renewal (Cheung and Rando, 2013). Reactivating the imprinted IGF2 gene expression or increasing mTOR activity in adult hematopoietic stem cells (HSCs) promote them to exit quiescence and proliferate (Chen et al., 2008; Venkatraman et al., 2013). IGF2 also stimulates mouse adult neural stem cell and intestinal stem cells reactivation (Ferron et al., 2015; Ziegler et al., 2015; Ziegler et al., 2019). In *Drosophila*, neural stem cells are reactivated in response to dietary amino acids and this has been attributed to insulin produced in glia cells (Chell and Brand, 2010; Sousa-Nunes et al., 2011; Huang and Wang, 2018). Elevated PI3 kinase-Akt-Tor activity stimulates reactivation of adult stem cells in *Drosophila* and mice (Chell and Brand, 2010; Sousa-Nunes et al., 2011; Gil-Ranedo et al., 2019; Hemmati et al., 2019). Although the role of the insulin/IGF-PI3 kinase-Akt-Tor signaling pathway has been demonstrated in many cell types and across a wide range of species (Chen et al., 2009; Ito and Suda, 2014; Ziegler et al., 2015), the mechanisms underlying its cell type-and cell state-specific activation are poorly understood. The similarities among mammalian, flies, and fish cells imply evolutionarily conserved and general mechanisms at work. Future studies are needed to determine whether the human PAPP-A/A2-mediated IGFBP proteolytic cleavage plays similar roles in the reactivation of HSCs and/or other quiescent cells.

## Materials and methods

### Experimental animals

Zebrafish were raised and crossed according to the standard husbandry guidelines (Westerfield, 2000). Embryos and larvae were raised at 28.5 C in the standard E3 embryo medium (Westerfield, 2000) unless stated otherwise. Two additional embryos media containing 0.2 mM [Ca^2+^] and 0.001 mM [Ca^2+^] (referred to as normal and low [Ca^2+^] medium, respectively) were prepared as previously reported (Dai et al., 2014) and used. 0.003% (w/v) N-phenylthiourea (PTU) was added to the media to inhibit pigmentation. *Tg(igfbp5a:GFP)* fish were generated in a previous study (Liu et al., 2017). The *pappaa*^*p170/+*^ fish were obtained from the Marc Wolman lab. All experiments were conducted in accordance with the guidelines approved by the Institutional Committee on the Use and Care of Animals, University of Michigan.

### Chemicals and reagents

Chemical and molecular biology reagents were purchased from Fisher Scientific (Pittsburgh, PA, USA) unless stated otherwise. BMS-754807 was purchased from JiHe Pharmaceutica (Beijing, China) and NBI-31772 from Tocris Bioscience (Ellisville, MO, USA). Liberase TM, Alizarin Red S, ZnCl_2_, and batimastat were purchased from Sigma (St. Louis, MO, USA). MK2206 was purchased from ChemieTek (Indianapolis, IN). Rapamycin was purchased from Calbiochem (Gibbstown, NJ). Antibodies (phospho-Akt, phospho-S6, and phospho-Erk) were purchased from Cell Signaling Technology (Danvers, MA, USA). Restriction enzymes were purchased from New England BioLabs (Ipswich, MA, USA). Primers, cell culture media, antibiotics, TRIzol, M-MLV reverse transcriptase, Alexa Fluor 488 Tyramide SuperBoost Kit, and Tetramethylrhodamine (TMRM) were purchased from Life Technologies (Carlsbad, CA, USA). Anti-Digoxigenin-POD and Anti-Digoxigenin-AP antibodies were purchased from Roche (Basel, Switzerland).

### Fluorescence activated cell sorting (FACS) and qPCR

FACS was performed to isolate NaR cells from *Tg*(*igfbp5a*:*GFP*) fish as described (Liu et al., 2017). Briefly, *Tg*(*igfbp5a*:*GFP*) larvae were raised in E3 medium to 3 days post fertilization (dpf) and transferred to normal or low [Ca^2+^] embryo media for 18 h. Single cell suspension was made using Liberase TM (0.28 Wünsch units/ml in HBSS). EGFP-positive cells were sorted using FACSAria Cell Sorter (BD Biosciences, Franklin Lakes, NJ). Total RNA was isolated using TRIzol LS reagent. Total RNA was reverse transcribed using SuperScript III Reverse Transcriptase. RNaseOUT^TM^ Recombinant Ribonuclease Inhibitor was added into the reverse transcript system to protect RNA from degradation. M-MLV Reverse Transcriptase was used for reverse transcription. qPCR was carried out using SYBR Green (Bio-Rad, Hercules, CA) on a StepONE PLUS real-time thermocycler (Applied Biosystems, Foster City, CA). Primers for qPCR were listed in Table S1.

### Genotyping

dCAPS assay and HRMA methods were used to genotype *pappaa* mutant. The dCAPS assay was performed following a published method (Wolman et al., 2015). Briefly, the region containing the *papp-aa* mutation was amplified with the dCAPS primers (Table S1), then followed by Mse1 digestion. To genotype pappaa mutant fish in a large number, HRMA were performed as previously reported (Liu et al., 2018) using the pappaa-HRMA primers (Table S1).

### Whole-mount *in situ* hybridization and immunostaining

A DNA fragment encoding part of Papp-aa sequence was amplified using primers shown in Table S1. The PCR product was cloned in pGEM-T easy plasmid. The Digoxygenin-UTP labeled sense and antisense riboprobes were synthesized as previously reported (Wang et al., 2009). Zebrafish larvae were fixed in 4% paraformaldehyde, permeabilized in methanol, and analyzed by whole mount immunostaining or *in situ* hybridization analysis as described previously (Dai et al., 2014).

For double color *in situ* hybridization and immunostaining, *pappaa* mRNA signal was detected using anti-DIG-POD antibody (Roche), followed by Alexa 488 Tyramide Signal Amplification (Invitrogen). After *in situ* hybridization analysis, the stained larvae were washed in 1xPBST and incubated with GFP antibody overnight at 4 °C. They were then stained with a Cy3 conjugated Goat anti-Rabbit IgG antibody (Jackson ImmunoResearch, West Grove, PA). Images were acquired using a Nikon Eclipse E600 Fluorescence Microscope with PMCapture Pro 6 software.

### Plasmid and BAC constructs

The expression plasmids for Papp-aa, Papp-ab, Papp-a2, Igfbp5a, Igfbp5a, and Igfbp3 have been reported (Kjaer-Sorensen et al., 2013, 2014). In order to generate cleavage resistant mutants of Igfbp5a and Igfbp5b, the following residues were mutated to alanine individually or in combination: Igfbp5a K147 and K148, Igfbp5b K144 and K145. Mutations were introduced by site-directed mutagenesis using QuikChange (Stratagene). The aforementioned plasmids were used as a template, and the primers used are shown in Table S1. All constructs were verified by sequencing analysis. The BAC constructions were generated as reported (Liu et al., 2017).

Briefly, zebrafish Papp-aa and Pappaa (E501A) cDNA were released from pcDNA3.1mH(+)A plasmids and sub-cloned into pIRES2-mCherry plasmid using SacII and BamH1 restriction sites. The pappaa/pappaa (E501A)-IRES2-mCherry-KanR cassette DNAs were amplified by PCR using primers containing 50bp homologies to *igfbp5a*. The DNA cassettes were inserted into the *igfbp5a* BAC construct to replace the *igfbp5a* sequence from the start codon to the end of the first exon through homologous recombination. A similar cloning strategy was used to generate *BAC(igfbp5a:hSTC1-IRES-mCherry)* and *BAC(igfbp5a:hSTC2-IRES-mCherry)* constructs. The construction of *BAC(igfbp5a: myrAkt-mCherry)* was reported previously (Liu et al., 2018). All constructs used were confirmed by DNA sequencing at the University of Michigan DNA Sequencing Core Facility.

For Tol2 transposon-mediated genetic mosaic assay, the validated BAC DNA and Tol2 mRNA were mixed and injected into *Tg(igfbp5a:GFP)* embryos at 1-cell stage. The embryos were raised and subjected to the low [Ca^2+^] stress test as shown in Fig. 2A. Cells co-expressing mCherry and GFP were identified and the cell division was scored following a reported scoring system (Liu et al., 2018).

### Live imaging and microscopy

NaR cells were quantified as previously reported (Liu et al., 2017). To visualize NaR cells, progeny of *papp-aa^+/−^;Tg(igfbp5a:GFP)* fish intercrosses were anesthetized with 0.63 mM tricaine. Live larvae were embedded in 0.3% low melting point agarose containing 0.63 mM tricaine and placed in a chamber in which the bottom was sealed with 0.16-0.19 mm slide. The solidarized agarose was then immersed in 1 ml normal [Ca^2+^] medium. The bright-field and GFP fluorescent images were acquired using Leica TCS SP8 confocal microscope equipped with HCPL APO 93X/1.30 GLYC. After taking images, the normal [Ca^2+^] medium was replaced and washed with low [Ca^2+^] medium for 3 times. The samples were incubated in low [Ca^2+^] medium for 8 hours and imaged again. LAS X and Image J were used for image analysis.

### Morphology analysis

Body length, defined as the curvilinear distance from the head to the end of caudal tail, was measured as reported (Liu et al., 2018). Somite number and head-trunk angles were measured manually as reported (Kamei et al., 2011). Alizarin red staining were performed as previously described (Liu et al., 2018). Images were captured with a stereomicroscope (Leica MZ16F, Leica, Wetzlar, Germany) equipped with a QImaging QICAM camera (QImaging, Surrey, BC, Canada).

### Drug treatment

Fish were treated with BMS-754807, ZnCl2, batimastat, and other drugs as reported previously (Liu et al., 2017). Drugs used in this study were dissolved in DMSO and further diluted in water. ZnCl_2_ which was dissolved in distilled water. Drug treatment was started at 3dpf unless stated otherwise. The drug solution was changed daily. The samples were collected for immunostaining after 24 hours treatment or for *in situ* hybridization after 48 hours treatment.

### Cell culture

Human embryonic kidney (HEK) 293 cells were cultured in DMEM (HEK 293) supplemented with 10% FBS, penicillin, and streptomycin in a humidified-air atmosphere incubator containing 5% CO_2_. Conditioned media were prepared as previously reported (Duan and Clemmons, 1998).

### Proteolytic assays and Western blotting

For *in vitro* proteinase assay, conditioned media were harvested from HEK293 cells transfected with zebrafish Igfbp5a, Igfbp5b, Igfbp3, Papp-aa, Papp-ab, Pappa2, and Papp-aa E501A plasmid alone or together. Conditioned media from MOCK transfected cells were used as controls. The media were concentrated and used in proteinase assay following published conditions (Kjaer-Sorensen et al. 2014). The proteinase assay was performed as described (Kjaer-Sorensen et al. 2014). Briefly, conditioned media collected from HEK293 cells transiently transfected with Papp-aa or Papp-ab were mixed 1:1 with media containing either Igfbp5a, Igfbp5b or Igfbp3 + 100nM IGF-II and incubated at 28°C. The reactions were stopped at different time points by adding 4x NuPAGE LDS sample buffer containing 75mM EDTA and 4mM PMSF. Next, proteins were separated by non-reducing SDS-PAGE. Subsequently, proteins were visualized by Western blotting as previously described (Boldt et al, 2001) using monoclonal anti c-myc antibody 9E10 (Thermo Fisher) 1:1000 and HRP-conjugated rabbit anti-mouse (P0260, DAKO) 1:2000.

### Statistical analysis

Values are shown as mean ± standard error of the mean (SEM). Statistical significance between experimental groups was determined using unpaired two-tailed t test, one-way ANOVA followed by Tukey’s multiple comparison test, Logrank test, or Chi-square test. Statistical significances were accepted at *P* < 0.05 or greater.

## Acknowledgments

We thank Dr. Marc Wolman, University of Wisconsin, for sharing the *pappaa*^*p170+/−*^ fish line. CD wishes to thank Dr. Stephen C. Blacklow, Harvard University, for his helpful discussions on LNR structure and regulation.

---

**Supplemental Figure 1.**
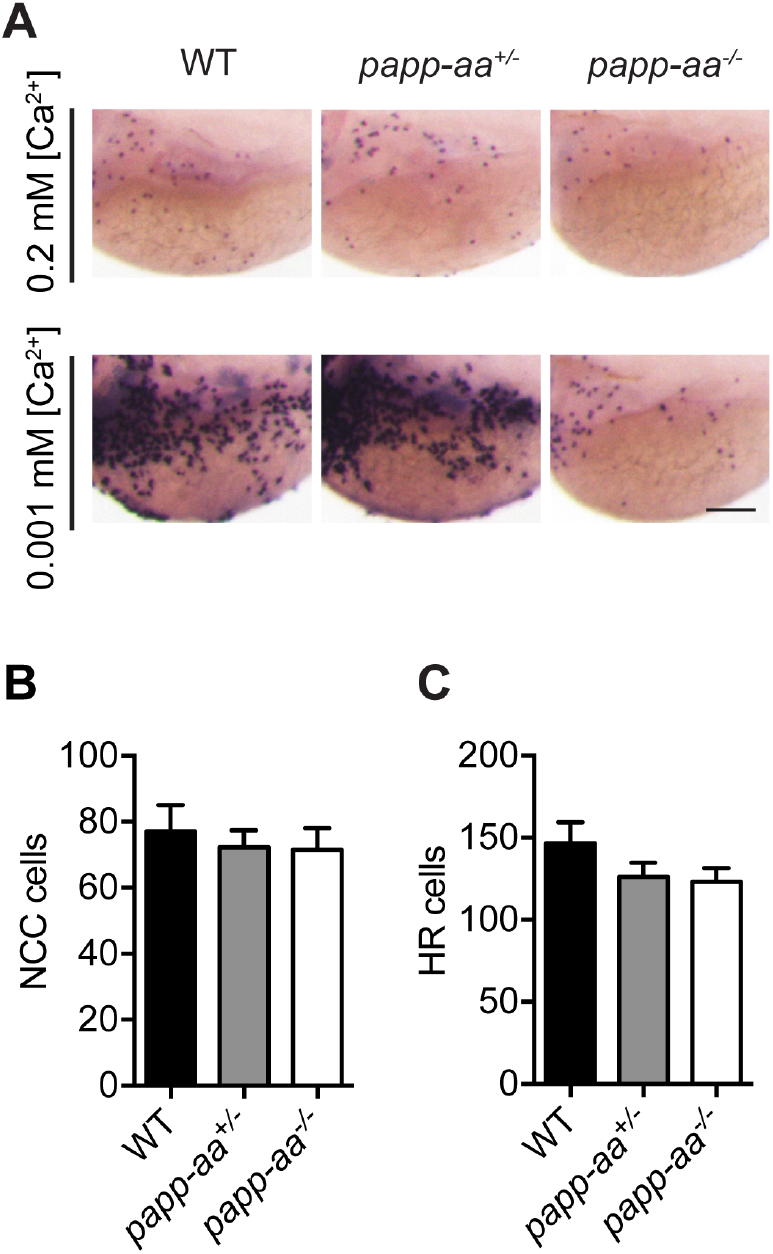
(**A**) Progeny of *papp-aa*^*+/−*^ intercrosses were treated as described in Fig. 2A. NaR cells were visualized by *trpv6* mRNA mRNA *in situ* hybridization. Scale bar = 0.1 mm. (**B-C**) Progeny of *papp-aa*^*+/−*^ intercrosses were treated as described in (**A**). NCC cells (**B**) and HR cells (**C**) were labeled by *slc12a10.2* mRNA and *atp6v1al* mRNA *in situ* hybridization and quantified. Each larva was genotyped afterwards. n = 7~31 fish/group.

**Supplemental Figure 2.**
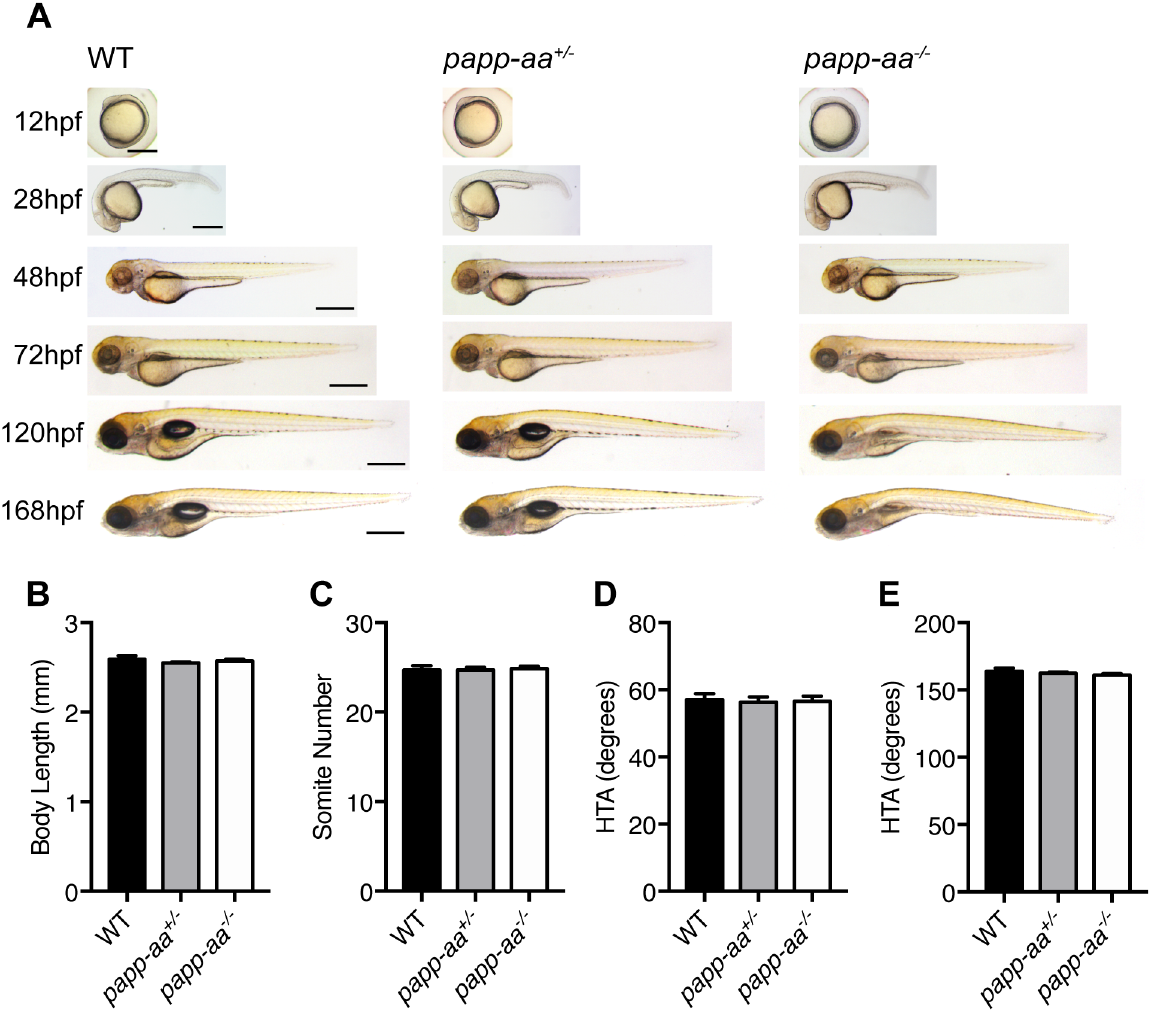
(**A**) Gross morphology of wild-type (WT), *papp-aa*^*+/−*^, and *papp-aa*^*−/−*^ fish at the indicated stages. Lateral views with anterior to the left. Scale bar = 0.5 mm. (**B-E**) Body length (**B**), somite number (**C**), and head-trunk angle (**D)** were measured at 24 hpf and head-trunk angle (**E**) at 3 dpf. Data shown are means ± SEM. n = 6~24 fish/group. No statistical significance was found using one-way ANOVA followed by Tukey’s multiple comparison test.

**Supplemental Figure 3.**
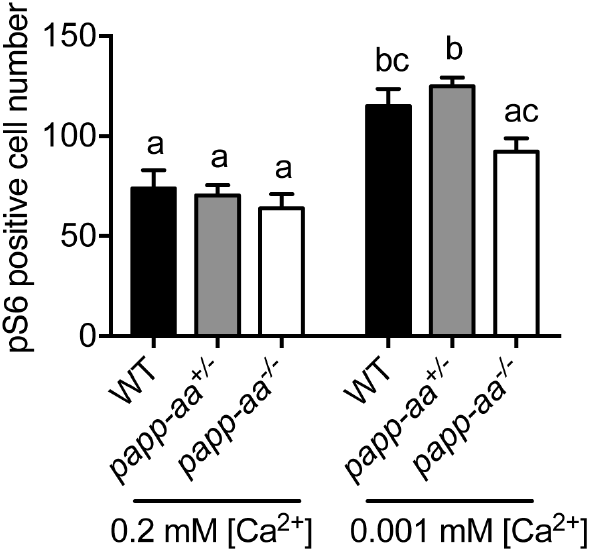
Loss of Papp-aa abolishes Ca^2+^ deficiency-induced Tor activity. Zebrafish of the indicated genotypes were transferred to normal or [Ca^2+^] medium at 3 dpf. One day later, they were fixed and stained for phospho-pS6. The number of phosphor-pS6-positive cells were quantified. n = 5~24.

**Supplemental Figure 4.**
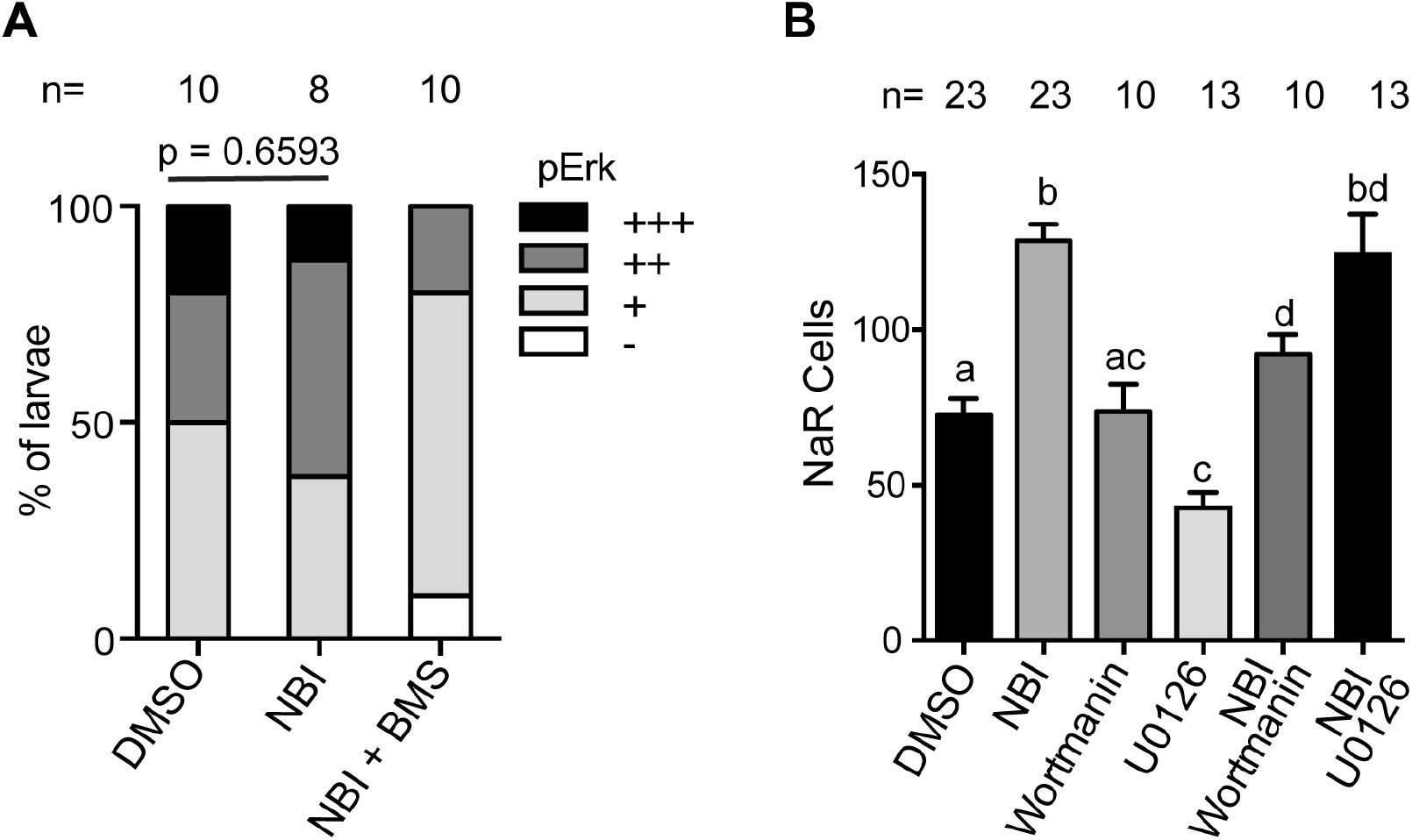
Disruption the IGF/Igfbp complex has no effect on pErk activity and Mek/Erk signaling is not critical. (**A**). Wild-type embryos were transferred to normal [Ca^2+^] medium containing 90 µM NBI-31772 with or without 0.3 µM BMS-754807. The fish were sampled at 4 dpf and subjected to immunostaining using a phosphorylated Erk antibody. The number of phospho-Erk-positive cells were scored following a published scoring system (Dai et al., 2014). Data shown are mean ± SEM, p values analyzed by Chi-square test for trend are shown. Total number of larvae is shown above the bar. (B) *Tg(igfbp5a:GFP)* larvae were treated with DMSO, NBI-31772 (NBI, 90 µM), Wortmannin (Wort, 0.06 µM), U0126 (2 µM), NBI-31772 (90 µM) plus Wortmannin (0.06 µM), or NBI-31772 (90 µM) plus U0126 (2 µM) from 3 to 5 dpf. NaR cell number was quantified and shown. n =10~23 fish/group.

## Notes

**Funding:** This work was supported by NSF grant IOS-1557850 and University of Michigan M-Cubed3 Project U064122 to CD and by a grant from DFF|FSS to CO. SL was supported by a fellowship from the China Oversea Scholarship Council. The funders had no role in study design, data collection and analysis, decision to publish, or preparation of the manuscript.

**Competing interests:** The authors declare that no competing interests exist.

